# The *Mycobacterium tuberculosis* long-chain fatty acid resistome reveals the universal stress protein TB15.3 as essential for infection

**DOI:** 10.1101/2025.04.14.648703

**Authors:** Alexandre J. Pinto, Marco Silva, Joana A. Santos, Brian C. VanderVen, Dirk Schnappinger, Sabine Ehrt, Tiago Beites

## Abstract

The human pathogen *Mycobacterium tuberculosis* (Mtb) thrives in lipid-rich microenvironments. Long-chain fatty acids (LCFA) are some of the most abundant lipids encountered by Mtb during infection. Mtb has evolved to utilize LCFA as a preferred carbon source, however, LCFA are also known to be potent antimicrobials. Mtb must therefore employ mechanisms to utilize LCFA as a carbon source while avoiding its bactericidal properties. We used transposon sequencing (TnSeq) to define a Mtb LCFA resistome, and found it to be composed by 38 genes. Surprisingly, Mtb requires a diverse set of metabolic pathways to avoid LCFA toxicity, indicating pleiotropic effects of LCFA in Mtb metabolism. As a functional follow-up on the TnSeq screen, we investigated the function of the universal stress protein TB15.3 in LCFA metabolism and during infection. We show that TB15.3 acts as a “metabolic break” for LCFA uptake and catabolism, avoiding deleterious membrane hyperpolarization. This was associated with loss of viability in the chronic phase of infection in mice and in an in vitro caseum model. Our work highlights Mtb LCFA resistance mechanisms as an important adaptation to the host and a promising target space to be exploited for drug development.

## INTRODUCTION

Tuberculosis (TB), caused by *Mycobacterium tuberculosis* (Mtb), is a leading cause of morbidity and mortality worldwide. In 2023, it is estimated that 1.3 million people died, while over 10.8 million fell ill from the disease^1^. TB treatment regimens are long and require multiple drugs, which contribute to treatment failure and create opportunities for the emergence of drug resistances^1^. Novel strategies to eradicate infections can unveil unexplored drug target spaces that can ultimately contribute to new treatment options for TB.

TB lesions are notorious for lipid accumulation^2-4^, an outcome that is partially driven by the bacteria^5^. Within the granuloma, a multicellular structure that constitutes the TB pathology hallmark, Mtb primarily inhabits macrophages that often acquire a lipid-laden phenotype or resides in the caseum, the necrotic center that is composed of an acellular matrix with high lipid content^6,7^. Lipid accumulation favors Mtb survival and induces a drug-tolerant phenotype, contributing to the need of long and multidrug treatment regimens^8,9^. To utilize fatty acids^10-13^ and cholesterol^14-16^ as carbon sources, Mtb relies on dedicated, complex uptake systems that require ATP. Mce1 facilitates the uptake of fatty acids^17^ while Mce4 mediates cholesterol uptake^14^. Interestingly, long-chain fatty acids (LCFA) are known to have antimicrobial properties^18^ through mechanisms that include cell membrane disruption^19^, interference with respiration^20^ or inhibition of essential membrane proteins^21^.

Although poorly understood, LCFA are known for a long time to be bactericidal to Mtb *in vitro*^22^. Analysis of caseum composition has revealed that LCFA are present at high concentrations (millimolar range)^9^. The dual nature of LCFA, both a carbon source and an antimicrobial, raises the hypothesis that Mtb evolved LCFA resistance mechanisms. Importantly, the inhibition of these mechanisms presents an opportunity to kill Mtb through intoxication with a preferred carbon source.

We have previously identified Mtb proteins required to resist LCFA. Inactivation of the respiratory enzyme Ndh-2^23^ or the *β*-oxidation related membrane oxidoreductase EtfD^24^ impacted both LCFA susceptibility and virulence, while inactivation of the adenylate cyclase MacE^25^ was shown to increase susceptibility to LCFA *in vitro*. Other bacteria present dedicated LCFA resistance mechanisms, like the *Staphylococcus aureus* enzyme FAME that esterifies LCFA with cholesterol to avoid toxicity^26,27^. Our previous work suggested that Mtb displays a diverse array of LCFA resistance mechanisms.

In this work, we aimed at defining Mtb’s LCFA resistome. We performed transposon sequencing (TnSeq) and identified 38 genes associated with increased LCFA susceptibility. Interestingly, the 38 genes encode proteins with diverse functions that span a wide array of metabolic pathways and cell processes. We then investigated the function of the screen’s top hit *tb15*.*3* (*rv1636*), which encodes a universal stress protein (USP) that binds cyclic AMP (cAMP) with high affinity^28,29^. Our data show that TB15.3 limits LCFA uptake and catabolism preventing deleterious membrane hyperpolarization. Importantly, we show that TB15.3 is essential to survive during the chronic phase of infection in mice, as well as in a caseum *in vitro* model.

## RESULTS

### The Mtb LCFA resistome

The dual nature of LCFA, as carbon source and antimicrobial, together with the previous identification of proteins needed to resist LCFA^23-25^ led us to hypothesize that Mtb employs multiple mechanisms to resist LCFA. To define Mtb’s LCFA resistome, we performed TnSeq (Fig. 1a), by comparing transposon libraries grown in Sauton’s with glucose/glycerol (fatty acid free - FAF) and the same medium supplemented with oleic acid (OA). The rationale behind this screen was to identify mutants that are sensitive to OA even if an alternative carbon source is provided.

**Fig. 1:**
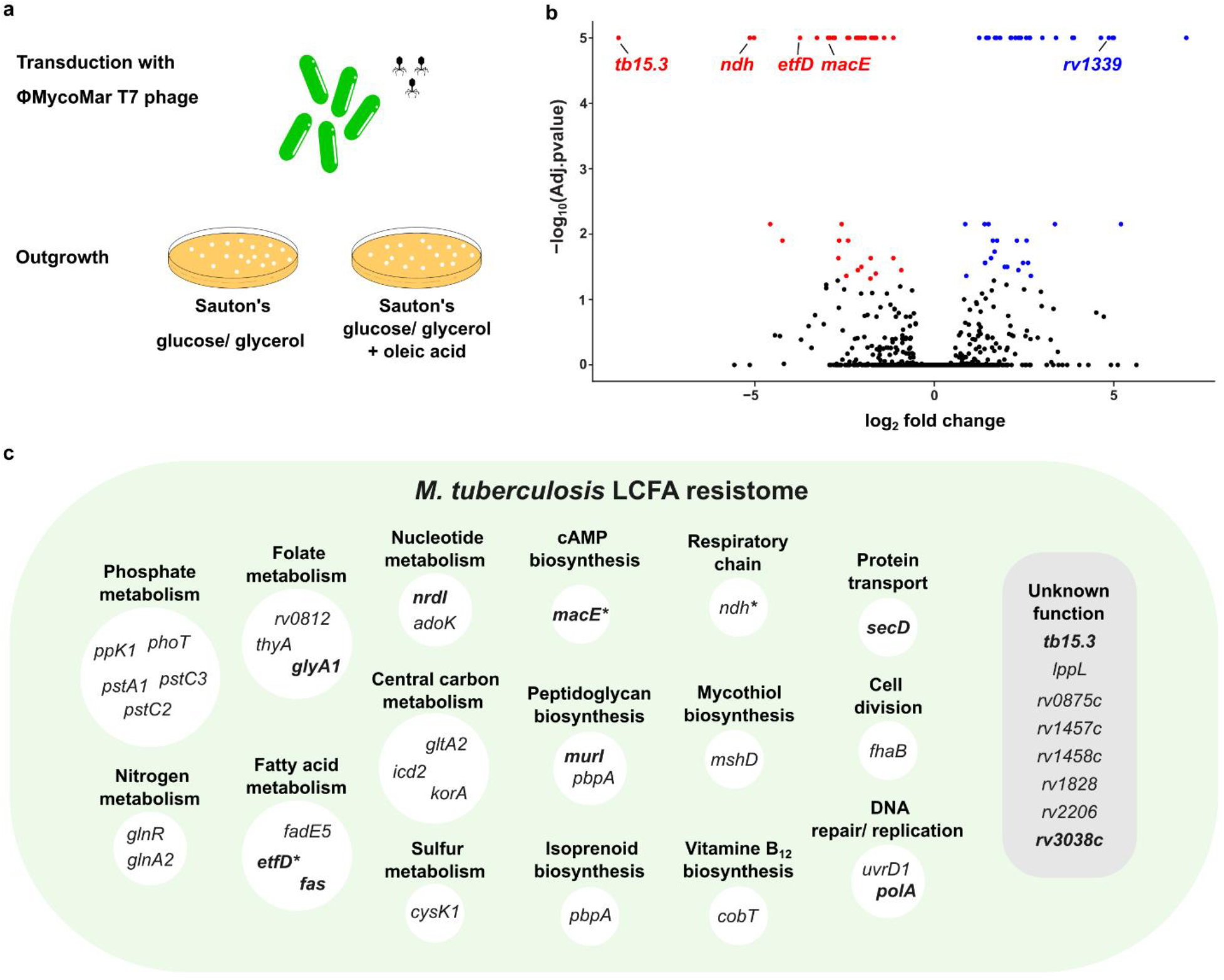
Transposon-sequencing (TnSeq). a) Cartoon representing the experimental setup. Transposon libraries were generated in Sauton’s glucose/ glycerol (fatty acid free – FAF libraries) and in Sauton’s glucose/ glycerol supplemented with oleic acid 200 *μ*M (OA libraries). b) Genes with statistically significant different transposon insertion counts: underrepresented (red) and overrepresented (blue) in OA libraries when compared with FAF libraries. Identified spots refer to genes previously associated with differential LCFA susceptibility (*ndh, etfD, macE* and *rv1339*), and the gene selected for functional follow-up (*tb15*.*3*). c) Functional categorization of genes predicted to participate in Mtb’s resistance to LCFA.

Using TRANSIT for statistical analysis^30^, we identified 38 genes with few or no insertions in the library exposed to OA when compared to FAF libraries (Fig. 1b and 1c; Supplementary data 1). Of these, 10 genes are predicted to be essential for growth in vitro (*etfD, glyA1, murI, polA, tb15*.*3, fas, secD, rv3038c, nrdI* and *macE*) and 11 genes are predicted to be associated with a growth defect (*rv0812, gltA2, rv1457c, rv1458c, rv1828, ndh, lppL, adoK, korA, thyA* and *ppK*)^31^. The previously described *ndh*^*23*^, *etfD*^*24*^, and *macE*^*25*^ were identified as part of the Mtb LCFA resistome, validating our screen. Genes involved in fatty acid metabolism (*etfD, fadE5* and *fas*) or central carbon metabolism (*gltA2, icd2* and *korA*) indicate that OA needs to be further metabolized to avoid toxicity. We also identified genes encoding enzymes that participate in pathways that are not directly connected with LCFA, such as folate metabolism (*rv0812, glyA1* and *thyA*), nitrogen metabolism (*glnR* and *glnA2*), nucleotide metabolism (*nrdI* and *adoK*), peptidoglycan biosynthesis (*murI* and *pbpA*) or phosphate metabolism (*ppk1, phoT, pstA1, pstC2* and *pstC3*) (Fig. 1c; Supplementary data 1). Finally, 8 genes of unknown function (*tb15*.*3, lppL, rv0875c, rv1457c, rv1458c, rv1828, rv2206* and *rv3038c*) were associated with increased OA susceptibility. This profile suggests that LCFA display pleiotropic metabolic interactions in Mtb metabolism and that multiple pathways need to be functional to avoid toxicity.

We also identified 46 genes that accumulated mutations in the OA-exposed library (Fig. 1b), from which the vast majority are predicted to be non-essential (40 genes). Among this gene set were the phosphodiesterase encoding gene *rv1339*, which has been implicated in LCFA metabolism and genes important for carbohydrate utilization (p.e *glpK*) and uptake (p.e *ppe51/pe19*). Other notable hits include genes encoding RelA (ppGpp synthesis), the two-component system PhoPR and cholesterol degradation enzymes.

This TnSeq screen identified the Mtb LCFA resistome, unveiling a largely unexplored drug target space.

### Universal stress protein TB15.3 is required to resist multiple LCFA

For a functional follow-up on the TnSeq screen, we focused on the top-hit *tb15*.*3* (*rv1636*) for the following reasons: i) the possibility of unveiling the physiological role of a protein of unknown function, ii) the predicted essentiality for in vitro growth^31,32^, iii) the considerable interest in TB15.3 function^28,29,33^, and iv) the possibility of characterizing a druggable target^34-36^. TB15.3 is a universal stress protein that binds cAMP with high affinity^28^. The ability to bind cAMP is necessary for the essential, yet undefined function of TB15.3^29^. However, our TnSeq screen predicts that *tb15*.*3* is not an essential gene, but rather a conditionally essential gene depending on the presence of LCFA in the environment. To test this prediction, we generated a *tb15*.*3* knockout strain using FAF medium for the outgrowth of transformants. In agreement with the TnSeq prediction, we successfully isolated a Mtb *tb15*.*3* knockout strain (Δ*tb15*.*3*) and confirmed its genetic identity through whole genome sequencing (WGS). Analysis for polymorphisms did not identify mutations that potentially impact growth (Supplementary Table 1). Additionally, we generated a Δ*tb15*.*3* complemented strain in which a copy of *tb15*.*3* was reintroduced.

We performed minimal inhibitory concentration (MIC) assays with fatty acids with different chain lengths and cholesterol to investigate if the observed hypersusceptibility of Δ*tb15*.*3* was OA specific. Mtb Δ*tb15*.*3* did not show altered susceptibility to short and medium-chain fatty acids (butyric acid, valeric acid, and octanoic acid), or to cholesterol when compared to the wild type and complemented strain (Fig. 2a, Supplementary Fig. 1). The profile with valeric acid and cholesterol also showed that this mutant is not hypersusceptible to the degradation product propionic acid, a metabolite that is bactericidal at high intracellular concentrations^37-39^. In contrast, Δ*tb15*.*3* displayed increased susceptibility to all LCFA tested (Fig. 2a). These LCFA, namely palmitic acid (PA), OA, and arachidonic acid (AA) were shown to be present in Mtb-generated lesions^9,13^, attesting to their physiological relevance. To understand if LCFA are bacteriostatic or bactericidal, we enumerated colony forming units (CFU) from the MIC plates. This revealed that all tested LCFA are bactericidal to Δ*tb15*.*3* at concentrations that support growth of wild type and complemented mutant (Fig. 2b).

**Fig. 2:**
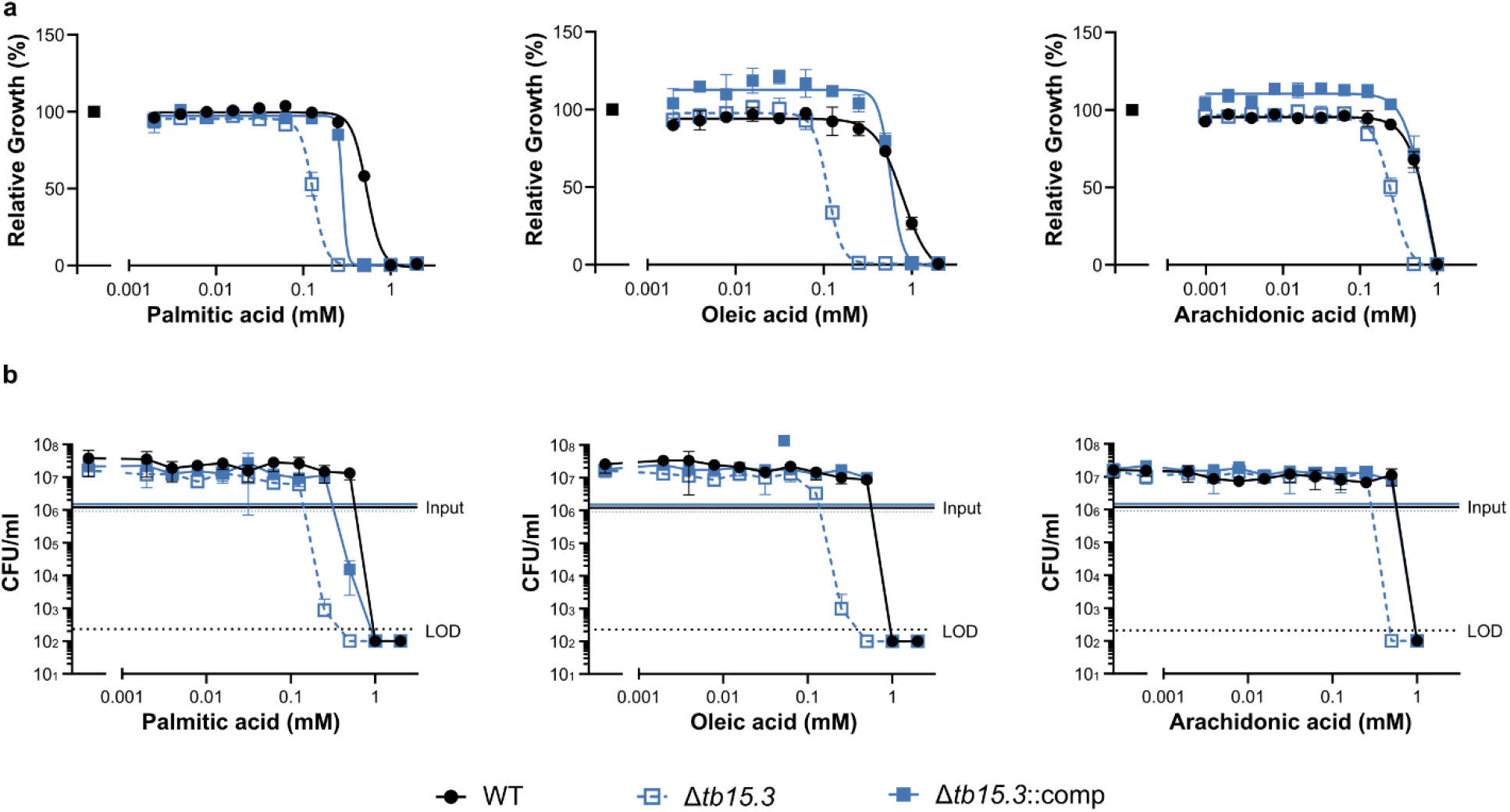
Long-chain fatty acids (LCFA) susceptibility profiles. a) Minimal inhibitory concentrations (MIC) of LCFA against Mtb wild type (WT), *tb15*.*3* knockout (Mtb Δ*tb15*.*3*) and complemented Mtb Δ*tb15*.*3* (Δ*tb15*.*3*::comp). Relative growth was calculated as OD_580nm_ values at day 14 of lipid over vehicle control. b) Bacterial viability estimation at the time of MIC OD_580nm_ measurements through CFU enumeration. LOD – limit of detection. Data are averages of three replicates and are representative of three independent experiments. Error bars correspond to standard deviation.

These data demonstrate that TB15.3 is essential to resist the antibacterial effects of LCFA.

### TB15.3 is required for a balanced LCFA uptake and catabolism

To understand how TB15.3 function integrates in the context of LCFA metabolism, we isolated 10 phenotypically confirmed OA resistant mutants in the Δ*tb15*.*3* genetic background (Supplementary Fig. 2). Polymorphism analysis revealed non-synonymous mutations, early stop codons and frameshift mutations in genes encoding components of the LCFA uptake transporter Mce1^17^ (Fig. 3a and 3b). This mutation profile strongly suggested that LCFA uptake is required for toxicity when TB15.3 is absent.

**Fig. 3:**
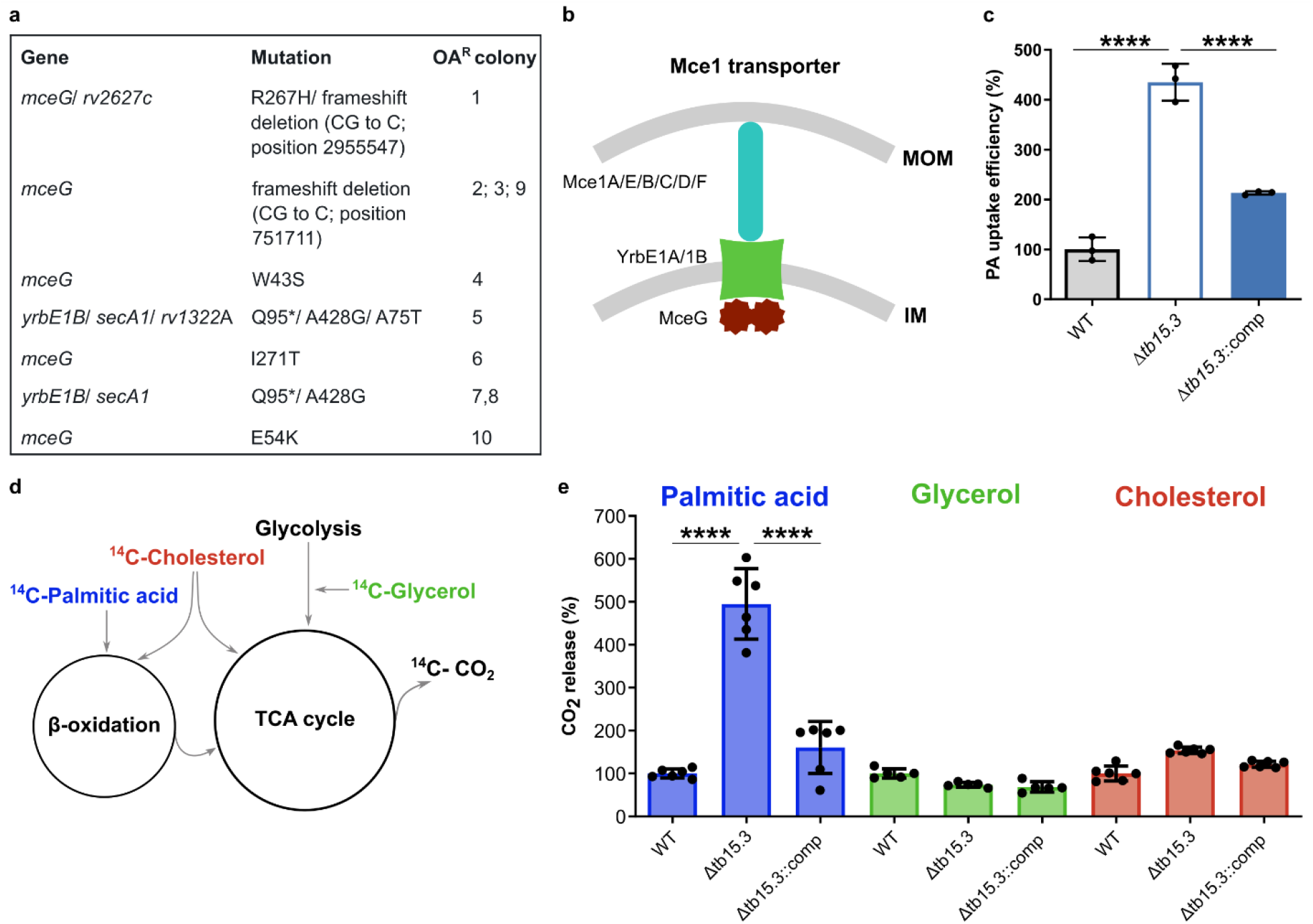
TB15.3 role in LCFA uptake and catabolism. a) Mutations detected in spontaneous oleic acid resistant (OA^R^) Mtb Δ*tb15*.*3* isolates. Depicted mutations represent the polymorphisms identified in OA^R^ colonies when compared with the parental strain (Mtb Δ*tb15*.*3*). * denotes early stop codon. b) Cartoon representing the structure of LCFA uptake complex Mce1. c) Efficiency of ^14^C-palmitic acid uptake in Mtb wild type (WT), *tb15*.*3* knockout (Mtb Δ*tb15*.*3*) and complemented Mtb Δ*tb15*.*3* (Δ*tb15*.*3*::comp). Data are averages of three replicates and are representative of two independent experiments. Error bars correspond to standard deviation. d) Cartoon representing Mtb carbon catabolic pathways. e) ^14^CO_2_ release assays Mtb wild type (WT), *tb15*.*3* knockout (Mtb Δ*tb15*.*3*) and complemented Mtb Δ*tb15*.*3* (Δ*tb15*.*3*::comp) grown in the presence of ^14^C-palmitic acid, ^14^C-glycerol and ^14^C-cholesterol. Data are averages of three replicates and are representative of two independent experiments. Error bars correspond to standard deviation. Statistical significance was assessed by one-way ANOVA followed by post hoc test (Tukey test; GraphPad Prism). \*\*\*\**P < 0*.*0001*

This led us to hypothesize that TB15.3 function might be associated with LCFA uptake. To test this, we measured the ^14^C-PA uptake rate of wild type, Δ*tb15*.*3* and complemented strains and found that the absence of TB15.3 leads to a significant increase of ^14^C-PA uptake, when compared to wild type and complemented strains (Fig. 3c). Next, we sought to investigate if this increase in PA uptake leads to increased LCFA degradation through *β*-oxidation/TCA cycle. We grew wild type, Δ*tb15*.*3* and complemented strains in medium with ^14^C-PA and measured the release of ^14^C-CO_2_ as a proxy for catabolic activity. The obtained profiles showed that Δ*tb15*.*3* displayed a 3-fold increase in ^14^C-CO_2_ release compared to wild type and complemented strains. To understand if these phenotypes were specific to LCFA, we performed the same experiment with carbon sources that are not toxic to Δ*tb15*.*3*, namely ^14^C-glycerol and _14_C-cholesterol. With these carbon sources the Δ*tb15*.*3* ^14^C-CO_2_ release profiles were similar to those of the wild type and complemented strains.

These results show that TB15.3 acts as a “metabolic brake” for uptake and catabolism of LCFA.

### Increased LCFA uptake and catabolism interfere with oxidative phosphorylation

To further understand the physiological role of TB15.3, and to test if this protein contributes to intrinsic drug resistance, we followed a chemical-genetics approach and determined the MIC profiles of antibiotics targeting different cellular functions: oxidative phosphorylation (OxPhos; DDD00853663, clofazimine, Q203 and bedaquiline), arabinogalactan biosynthesis (ethambutol), mycolic acid biosynthesis (isoniazid), transcription (rifampicin), translation (amikacin and linezolid), peptidoglycan biosynthesis (vancomycin) and DNA replication (moxifloxacin). MICs were determined in two different media: Sauton’s glucose/glycerol and Sauton’s glucose/glycerol supplemented with OA 100 *μ*M (subinhibitory concentration for Δ*tb15*.*3*). In Sauton’s glucose/glycerol, Δ*tb15*.*3* was similarly susceptible as wild type to all tested antibiotics (Supplementary Table 2). However, in the OA-supplemented media, Δ*tb15*.*3* showed increased susceptibility to all antibiotics targeting oxidative phosphorylation and no reproducible differential susceptibility to the other antibiotics (Fig. 4a and 4b, Supplementary Table 2). Thus, TB15.3 is an intrinsic resistance factor to antibiotics targeting OxPhos, and connected Δ*tb15*.*3* LCFA susceptibility to interference with oxidative phosphorylation.

**Fig. 4:**
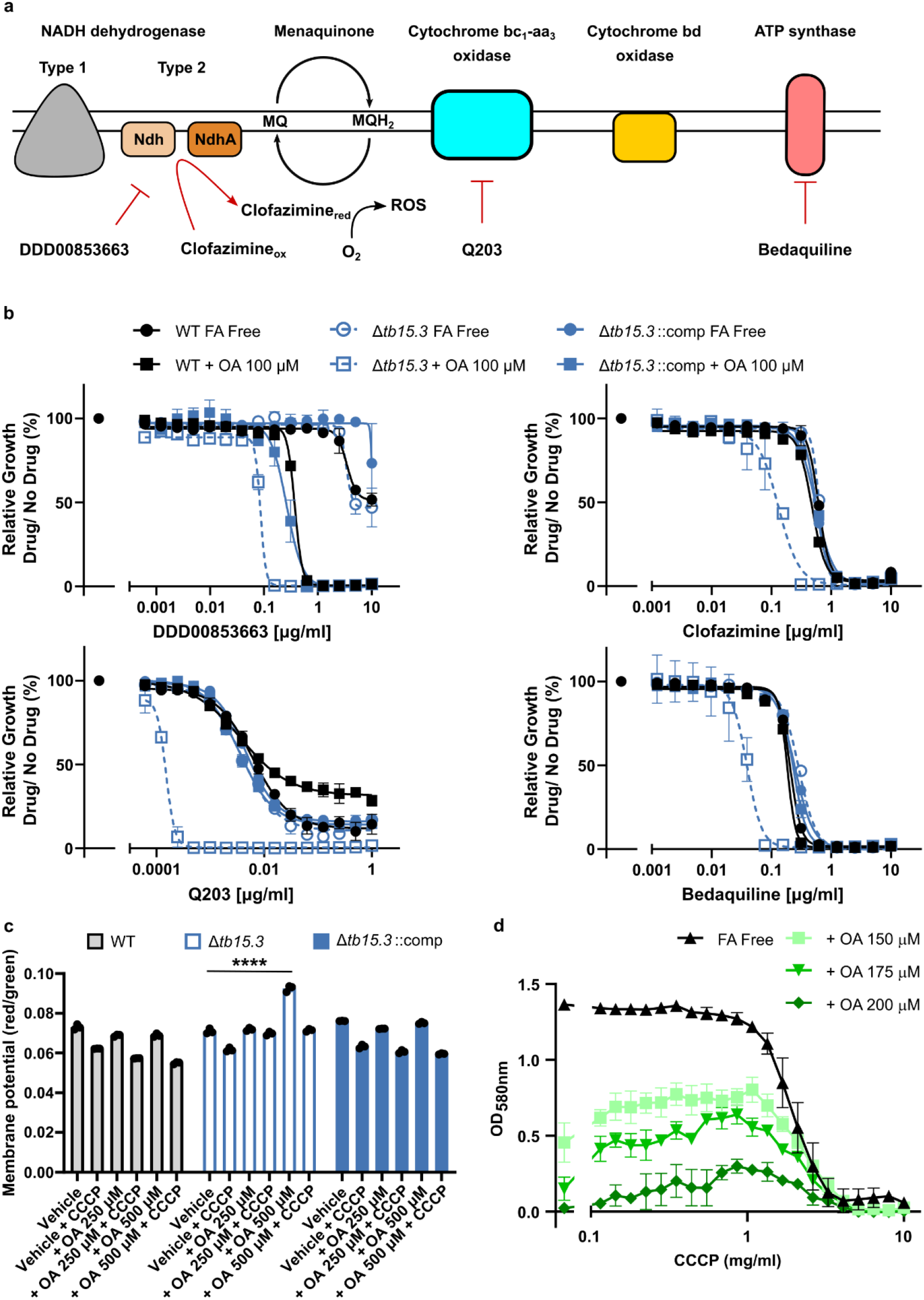
Chemical-genetics identify TB15.3 as an intrinsic resistance factor to oxidative phosphorylation inhibition. a) Cartoon representing Mtb’s oxidative phosphorylation and inhibitors mode of action. b) Susceptibility profiles of Mtb wild type (WT), *tb15*.*3* knockout (Mtb Δ*tb15*.*3*) and complemented Mtb Δ*tb15*.*3* (Δ*tb15*.*3*::comp) to Ndh-2 inhibitor DDD00853663, Ndh-2 engaging drug clofazimine, cytochrome bc1-aa3 oxidase inhibitor Q203 and ATP synthase inhibitor bedaquiline in modified Sauton’s fatty acid free (FA free) and the same medium supplemented with 100 *μ*M oleic acid (OA). Data are averages of three replicates and are representative of three independent experiments. Error bars correspond to standard deviation. c) Membrane potential in WT, Δ*tb15*.*3* and Δ*tb15*.*3*::comp treated with OA 250*μ*M and 500*μ*M or vehicle control for 24 hours. CCCP was used as a control for a depolarized membrane. Data are averages of three replicates and are representative of three independent experiments. Error bars correspond to standard deviation. Statistical significance was assessed by one-way ANOVA followed by post hoc test (Tukey test; GraphPad Prism). \*\*\*\**P < 0*.*0001*. d) Strain Δ*tb15*.*3* was treated with a concentration gradient of CCCP in different media: modified Sauton’s (FA Free) and in the same medium supplemented with different OA concentrations. OD_580nm_ measurements were taken after 14 days of culture. Data are averages of three replicates and are representative of three independent experiments. Error bars correspond to standard deviation.

Following up on this bioenergetics connection, we measured membrane potential, intracellular ATP levels and oxygen consumption rate (OCR), in response to treatment with OA. This revealed that the Δ*tb15*.*3* membrane became hyperpolarized in the presence of OA 500 *μ*M, while no effect was observed in the wild type and complemented strains (Fig. 4c). This was accompanied with a small burst in ATP levels in Δ*tb15*.*3* (Supplementary Fig. 3a). Given that the medium was at neutral pH, and thus the protonmotive force (PMF) is mainly driven by membrane potential, membrane hyperpolarization is expected to increase the activity of ATP synthase. OA had no impact on oxygen consumption in any of the strains (Supplementary Fig. 3b). To test if membrane hyperpolarization participates in LCFA toxicity in Δ*tb15*.*3*, we sought to rescue OA susceptibility with subinhibitory concentrations of the protonophore CCCP (Fig. 4d). Using a MIC setup in media with increasing concentrations of OA, we were able to identify a range of CCCP concentrations that allow Δ*tb15*.*3* to grow above OA MIC. Using a MIC setup in media with increasing concentrations of OA, we were able to identify a range of CCCP concentrations that increased resistance of Mtb Δ*tb15*.*3* to OA.

### TB15.3 is essential for Mtb survival during chronic mouse infection and in a caseum surrogate model

To understand if TB15.3 is required during infection, we infected C57BL/6J mice through aerosolization with approximately 100 CFU of wild-type, Δ*tb15*.*3* and complemented strain. We have followed the bacterial loads in lungs and spleens over a period of 70 days post-infection (dpi) through CFU enumeration. Results showed that, in lungs, Δ*tb15*.*3* was attenuated for growth and suffered from a survival defect during the chronic phase of infection (Fig. 5a). In contrast, wild type and complemented strain bacterial loads peaked at 28 dpi and stabilized until 70 dpi. Analysis of lung histopathology demonstrated that Δ*tb15*.*3* infected mice contained fewer and smaller pulmonary lesions resulting in an overall reduced lesion area than found in the mice infected with wild type or complemented strain (Fig. 5b and 5c). Regarding spleens, Δ*tb15*.*3* presented a similar tendency during chronic phase of infection, although at a lesser extent when compared to lungs indicating that the requirement for *tb15*.*3* is, at least partially, organ-specific.

**Fig. 5:**
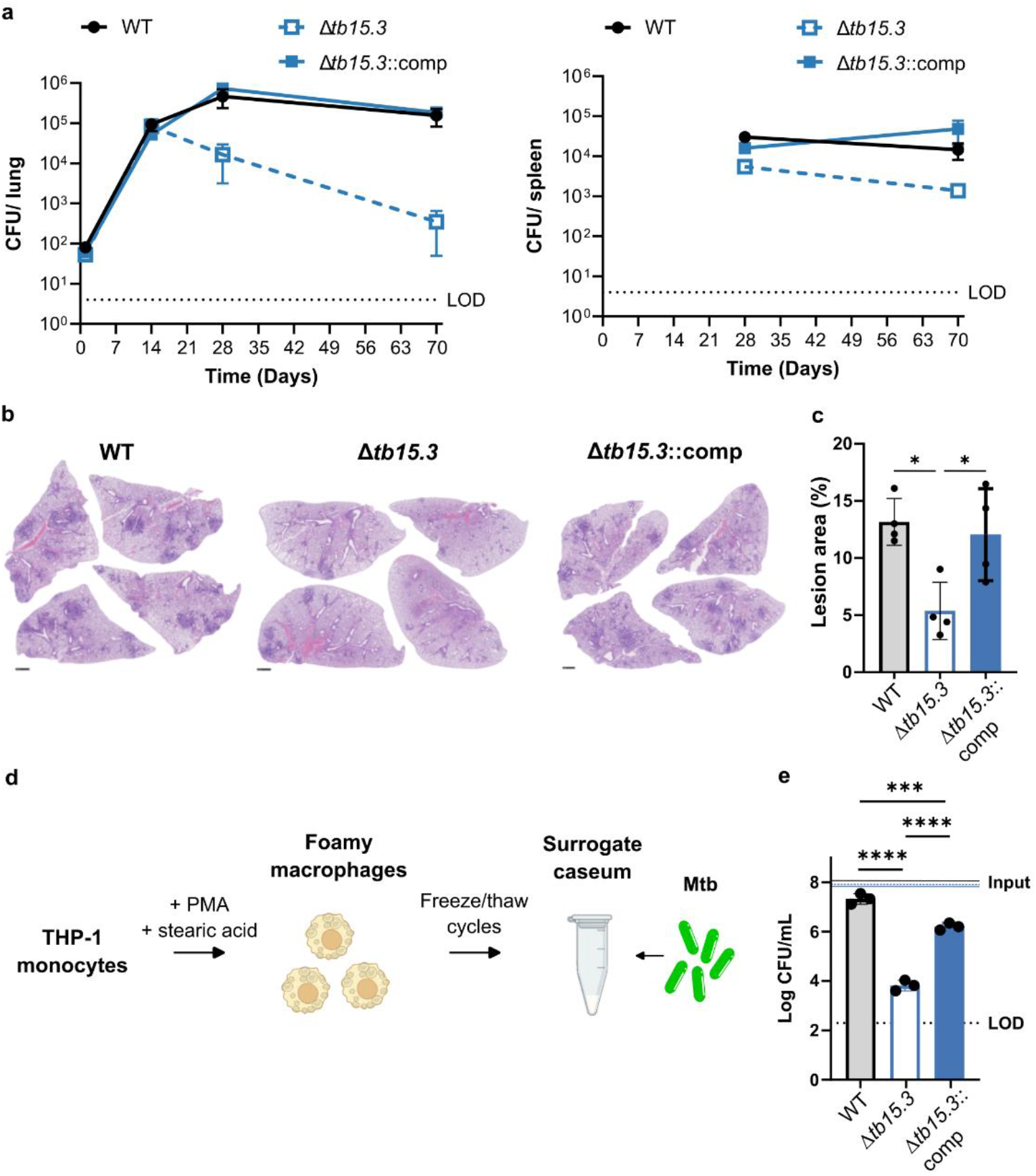
TB15.3 relevance for infection. a) Bacterial load over time of mice infected with Mtb wild type (WT), *tb15*.*3* knockout (Mtb Δ*tb15*.*3*) and complemented Mtb Δ*tb15*.*3* (Δ*tb15*.*3*::comp). LOD stands for limit of detection. Data are CFU averages from four mice and are representative of two independent experiments. Error bars correspond to standard deviation. b) H&E staining of lung histology sections. c) Quantification of lesion area based on histology represented in b). d) Cartoon representing caseum surrogate generation. e) Survival of WT, Δ*tb15*.*3* and Δ*tb15*.*3*::comp in caseum surrogate after 7 days estimated by CFU enumeration. LOD stands for limit of detection. Data are averages of three replicates and are representative of two independent experiments. Error bars correspond to standard deviation. Statistical significance was assessed by one-way ANOVA followed by post hoc test (Tukey test; GraphPad Prism). *** *P<0*.*001*; \*\*\*\**P < 0*.*0001*.

Mtb infected C57BL/6J mice do not generate necrotic lesions, and thus do not recapitulate important features of human TB lesions like caseum, the lipid-laden acellular matrix in the center of necrotic granulomas. Importantly, bacteria in caseum acquire a drug tolerant phenotype, which contributes to the need of long and multidrug treatment regimens^8^. Given that caseum contains high concentrations of LCFA^9^, we hypothesized that TB15.3 is required to survive in such an environment. To test this, we determined viability of wild-type, Δ*tb15*.*3* and complemented strains in a caseum in vitro model through CFU enumeration (Fig. 5c). As previously described, wild type Mtb entered a slow- to non-replicating state^9^. In contrast, Δ*tb15*.*3* lost viability (approximately 3-logs) when incubated in caseum, and this phenotype was complemented (Fig. 5d).

Altogether, these results show that TB15.3 is essential for optimal growth in lungs and for survival during chronic infection in mice, as well as in the caseum in vitro model.

## DISCUSSION

Throughout the co-evolution with the human host, Mtb developed a wide array of strategies to evade host defenses, adapt to heterogeneous niches and transmit to new hosts^40^. A common characteristic among host microenvironments faced by Mtb during infection is the accumulation of lipids^7^. Mtb thus evolved to utilize lipids, such as cholesterol and LCFA, as main carbon sources^41^. However, LCFA are also potent bactericidal molecules^18^. This provokes the hypothesis that LCFA resistance is an important host adaptation for Mtb. In fact, LCFA resistance mechanisms have been described for other bacteria^18^. Moreover, this dual nature of LCFA opens the possibility of eradicating Mtb infections through intoxication with a major host-derived carbon source.

We resorted to TnSeq to define how Mtb resists LCFA. This screen identified 38 mutants that were depleted in the presence OA when compared with the FAF libraries, thus predicting a role for the interrupted genes in LCFA resistance. The presence of genes previously identified, such as *ndh*^*23*^, *etfD*^*24*^ and *macE*^*25*^ among the 38 hits validated our TnSeq screen. LCFA resistance genes were enriched for genes classified as essential or associated with a growth defect, which can be explained by the fact that gene essentiality classifications in Mtb are generally based on genetic screens performed in media supplemented with LCFA^31,42,43^. One exception is the TnSeq reported by Minato, et al^44^, where libraries were generated in minimal medium and in which most of the genes identified in our screen as OA-sensitive were classified as non-essential. Our TnSeq re-classifies these predicted essential genes as conditionally essential depending on the presence of LCFA. Another unexpected aspect of our TnSeq data is the identification of genes encoding enzymes that participate in a diverse array of cell processes and metabolic pathways. The presence of genes related to fatty acid degradation and biosynthesis, as well as central carbon metabolism showed that the ability to metabolize LCFA once internalized is an important step towards LCFA resistance. The absence of lipid storage associated genes, such as *tgs1* (triglyceride synthase)^11^, suggest that this pathway does not participate in LCFA resistance in Mtb. Among the pathways lacking a direct connection with LCFA, phosphate metabolism was particularly enriched for genes associated with LCFA resistance. Polyphosphate (synthesized by Ppk1) was recently shown to play an important role in Mtb lipid biosynthesis, which may link phosphate metabolism with LCFA resistance^45^. Of note, it is possible that some of the 38 genes are exclusively associated with OA sensitivity. The precise definition of the LCFA resistome will, thus, require additional genetic screens for sensitivity to a broad range of LCFA. Nevertheless, all the proteins characterized so far revealed sensitivity to several different LCFA.

As a functional follow-up on the TnSeq, we characterized the protein of unknown function TB15.3. Half of Mtb proteins do not have a described function, which greatly limits our understanding of this pathogen’s biology^46^. Thus, we took this opportunity to functionalize a protein that has attracted considerable attention due to its predicted essentiality. The Mtb genome harbors 10 genes encoding USPs^46^. From these, Rv2623 was shown to be necessary for the establishment of a chronic infection^47,48^. Deletion of *rv2623* resulted in hypervirulence, as the mutant strain continued to grow after adaptive immunity activation^47,48^. Regarding *tb15*.*3*, it is the only USP encoding gene predicted to be essential for growth in vitro^31,32^. TB15.3 consists of a single USP domain and previous biochemical characterization showed that it binds cAMP with high affinity^28^. Importantly, cAMP binding was shown to be necessary for TB15.3 activity^29^. Our TnSeq screen predicted that *tb15*.*3* is conditionally essential depending on the presence of LCFA, which was confirmed by the successful isolation of a *tb15*.*3* knockout.

Characterization of Δ*tb15*.*3* showed that different LCFA are bactericidal to Δ*tb15*.*3* at concentrations permissive for growth of wild type and complemented strain, demonstrating that TB15.3 is a *bona fide* LCFA resistance factor. To understand the role of TB15.3 in LCFA metabolism, we raised spontaneous OA-resistant mutants in the genetic background of Δ*tb15*.*3*. All phenotypically confirmed OA-resistant mutants contained mutations in genes encoding the LCFA-uptake transporter Mce1^17^, strongly suggesting that LCFA need to be internalized to be toxic. This is in contrast with the LCFA antimicrobial activity against other bacteria, which generally encompasses direct interactions with the cell envelope^20^. The absence of TB15.3 was shown to lead to an increase in the uptake of LCFA, which correlates well with Mce1 inactivation preventing LCFA toxicity. Our results further show that Δ*tb15*.*3* degrades LCFA at a faster rate than the wild type and complemented strains, which may be due to increased LCFA availability and/or TB15.3 modulating the activity of *β*-oxidation/TCA cycle enzymes. Interestingly, in *M. smegmatis* the absence of the TB15.3 homologue increased glycerol degradation^33^. These data demonstrate that TB15.3 functions as a “metabolic brake” on Mtb LCFA uptake and catabolism.

How TB15.3 modulates LCFA uptake and catabolism requires further investigation. In *Escherichia coli*, the universal stress protein encoding gene *uspA* is repressed by the fatty acid degradation regulator FadR, although UspA function seems to be more related with adaptation to growth arrest^49^. Characterization of Mtb’s FadR homologue showed that it does not regulate *tb15*.*3* expression^50,51^. Moreover, analysis of Mtb’s transcriptome grown in a lipid rich environment did not reveal upregulation of *tb15*.*3* expression^52^. Interestingly, when Mtb cAMP intracellular levels get depleted, an increase in LCFA uptake/catabolism associated with OA-sensitivity has been observed^25^, phenocopying what we observe in Δ*tb15*.*3*. Hence, given that a functional TB15.3 requires cAMP binding^29^, it is likely that cAMP intracellular levels regulate TB15.3 function. Regarding TB15.3 activity, two hypotheses have been proposed: 1) it may modulate the levels of free intracellular cAMP^28^ and 2) it may act through protein-protein interaction to modulate activity of interacting proteins from central carbon metabolism^33^. In the future, we will further characterize TB15.3 biochemically to test these hypotheses.

The presence of LCFA in the medium sensitized Δ*tb15*.*3* to OxPhos inhibition. That LCFA sensitizes Δ*tb15*.*3* to compounds targeting different OxPhos enzymes, suggests a pleiotropic effect rather than the inhibition of a specific enzyme. The increased sensitivity to OxPhos inhibitors can, thus, be due 1) to excess reducing equivalents coming from LCFA degradation that have to be oxidized by respiration enzymes, putting pressure on the electron transport chain, or 2) the direct inhibition of respiration by LCFA, as observed in other bacteria^53^. We did not observe inhibition of oxygen consumption or a decrease in intracellular ATP levels in Δ*tb15*.*3* treated with OA, which argues in favor for the need of functional OxPhos to deal with the reducing equivalents generated by LCFA degradation. This profile linked OA toxicity with bioenergetics, and, in fact, we demonstrated that membrane hyperpolarization contributes to toxicity in Δ*tb15*.*3*. This is in contrast with other bacteria where LCFA leads to a depolarized membrane^54^. The chemical-genetics data also showed that targeting TB15.3 can potentiate the action of drugs in clinical trials (Q203) and in clinical use (bedaquiline and clofazimine) for TB treatment.

We further showed that TB15.3 is necessary for optimal growth during acute infection and essential for survival during chronic infection in mice. Moreover, it is also essential for survival in a caseum *in vitro* model. USP were first described and have been consistently described as proteins required to endure stress^55^. To survive the multiple stresses during chronic infection induced by adaptive immunity, Mtb becomes less metabolically active^41^. Thus, it is possible that the loss of an important “metabolic brake” like TB15.3 counteracts the need of a slower metabolism, contributing to loss of viability. In the case of caseum, the bacteria are embedded in an environment with readily available LCFA at concentrations in the millimolar range^9^, which strongly suggests that in this case Δ*tb15*.*3* loses viability directly through LCFA toxicity.

Our study highlights LCFA resistance as an important adaptation of Mtb to the host. The functional analysis of the LCFA resistance factor TB15.3 revealed a “metabolic brake” necessary to keep LCFA uptake and catabolism in homeostasis, which is essential for survival during chronic infection and in caseum. The identification of compounds that bind TB15.3 suggests that it may be a druggable target^34,36^. Importantly, given that Mtb inhabits lipid-laden environments during infection, the LCFA resistome constitutes an attractive target space for drug development.

## METHODS

### Growth conditions

Escherichia coli was used as a host for cloning and was cultured in LB medium at 37 °C. *M. tuberculosis* was cultured at 37 °C in a modified Sauton’s minimal medium (0.05 % potassium dihydrogen phosphate, 0.05 % magnesium sulfate heptahydrate, 0.2 % citric acid, 0.005 % ferric ammonium citrate, and 0.0001 % zinc sulfate) supplemented with 0.05 % tyloxapol, 0.4 % glucose, 0.2 % glycerol, and ADNaCl with fatty acid free BSA (Roche). Modified Sauton’s solid medium contained 1.5 % bactoagar (BD) and glycerol at a higher concentration (0.5 %). OA, PA and AA were dissolved in a tyloxapol:ethanol (1:1) solution and added directly to cultures. Antibiotics were used at the following final concentrations: carbenicillin 100 μg/ml, hygromycin 50 μg/ml and kanamycin 50 μg/ml.

### Mutant construction

Deletion of *tb15*.*3* native locus was achieved through recombineering. A DNA fragment with the hygromycin resistant gene (hygR) flanked by 500 bp upstream and downstream of *tb15*.*3* was synthesized (GeneScript) and transformed in Mtb H37Rv with a plasmid expressing the recombinase RecET. Transformants were selected in fatty acid free modified Sauton’s supplemented with hygromycin. The recET plasmid was counterselected by growing transformants in modified Sauton’s supplemented with sucrose 10 % (w/v). To generate the complemented strain, we have cloned *tb15*.*3* in an integrative plasmid (att-L5 site) with a kanamycin resistant cassette and the constitutive promoter (pMCK-phsp60-*tb15*.*3*) and then transformed it in *tb15*.*3* knockout.

### Transposon sequencing

Himar1 transposon mutant libraries were constructed in Mtb H37Rv by transduction with the ΦMycoMar T7 phage^56^, and grown in either modified Sauton’s solid medium or modified Sauton’s solid medium supplemented with OA 200 *μ*M. Plates were incubated at 37 °C for 28 days and then colonies were scraped and pooled. Genomic DNA was extracted, fragmented, the transposon–chromosome junctions were amplified and sent for sequencing to quantify the relative abundance of transposon insertions, as described previously described. Sequencing reads processing and statistical analysis of transposon insertion quantification were performed in the TRANSIT TnSeq analysis platform^30^.

### Minimal inhibitory and bactericidal concentration experiments

Strains were cultured modified Sauton’s medium until the mid-exponential phase (Optical density at 580nm - OD_580nm_ of 0.5-1), centrifuged (4500 rpm; 10 min), resuspended in fresh medium and centrifuged again (800 rpm; 10 min) to generate single bacterial suspensions. For LCFA MIC assays, we have made serial dilutions (two-fold) in 96-well plates and added bacteria to a final volume of 200 *μ*l and a final OD_580nm_ of 0.01. Plates were incubated at 37 °C for 14 days and then OD_580nm_ was registered. CFU enumeration from all wells was then performed to estimate MBC. Modified Sauton’s solid medium was used for outgrowth. For compounds MIC assays and CCCP rescue experiments, we have used 384-well plates. Compounds were dispensed as serial dilutions (two-fold) in a D300e Digital Dispenser (HP). Bacteria were then added to a final volume of 50 *μ*l and a final OD_580nm_ of 0.01. DMSO was normalized across wells to 1 % (vol/vol). Plates were incubated at 37 °C for 14 days and then OD_580nm_ was registered.

### Spontaneous mutant screen

The strain Δ*tb15*.*3* was grown in modified Sauton’s medium until stationary phase (21 days). Solid modified Sauton’s medium was supplemented with OA 500 μM, a condition that is not growth permissive to Δ*tb15*.*3*, to select OA-resistant spontaneous mutants in the Δ*tb15*.*3* genetic background. To this end, we inoculated the solid medium with different bacterial densities (10^7^, 10^8^ and 10^9^) and incubated the plates for 4 weeks. To assure that the bacteria were viable we also inoculated solid Sauton’s medium (fatty acid free). Colonies that grew in the presence of OA were picked and grown in liquid modified Sauton’s medium. To confirm the ability to grow in the presence of LCFA, we cultured WT, Δ*tb15*.*3*, complemented strain and spontaneous mutants in modified Sauton’s medium supplemented with OA 500 *μ*M and PA 250 *μ*M and measured OD_580nm_ after 21 days.

### Whole genome sequencing

Confirmation of *tb15*.*3* deletion and identification of genetic variants in spontaneous OA-resistant mutants were performed by WGS. Briefly, genomic DNA (150-200 ng) was sheared and HiSeq sequencing libraries were prepared using the KAPA Hyper Prep Kit (Roche). PCR amplification of the libraries was carried out for ten cycles. 5–10 × 10^6^ 50-bp paired-end reads were obtained for each sample on an Illumina HiSeq 2500 using the TruSeq SBS Kit v3 (Illumina). Post-run demultiplexing and adapter removal were performed and fastq files were inspected using fastqc^57^. Trimmed fastq files were then aligned to the reference genome (M. tuberculosis H37RvCO; NZ_CM001515.1) using bwa mem^58^. Bam files were sorted and merged using samtools^59^. Read groups were added and bam files de-duplicated using Picard tools and GATK best-practices were followed for SNP and indel detection^60^. Gene knockouts and cassette insertions were verified by direct comparison of reads spanning targeted sequences to plasmid maps and the genome sequence.

### Lipid uptake and catabolic rate measurements

LCFA uptake was quantified as previously described^17^. Mtb was cultured in 7H9 containing fatty acid-free albumin-dextrose supplemented with 0.01% glycerol and 0.05% tyloxapol (vented T-25 tissue culture flasks). Mid-logarithmic growth phase bacteria were concentrated in a spent medium to a final OD_600nm_ of 0.7. The bacterial cultures were immediately supplied 1.0 µCi of [1-^14^C]-PA (Perkin Elmer) and were incubated at 37 °C. Samples of 1.5 ml from the cultures were harvested at 5, 30, 60, and 120 min. Samples were washed three times with cold PBS containing 0.1% Triton X-100 and 0.1% fatty acid-free BSA to remove surface-bound radiolabel. Then bacteria were fixed in 4% paraformaldehyde (PFA) and ^14^C incorporation was quantified via scintillation counting. Radioactive counts at each time point were plotted and used for linear regression calculations to determine the rate of lipid uptake. The rates were normalized to wild type and expressed as uptake efficiency (%).

Catabolism was measured by the quantification of ^14^CO2 released from [1-^14^C]-PA, from [1-^14^C]-glycerol and [1-^14^C]-cholesterol (Perkin Elmer). Mtb was cultured in 7H9 containing fatty acid-free albumin-dextrose supplemented with 0.01 % glycerol and 0.05 % tyloxapol to the mid-logarithmic phase of growth (vented T-25 tissue culture flasks). Cultures were then concentrated in spent medium to an OD_600nm_ of 0.5 and 1.0 μCi of radiolabeled carbon sources was added to each flask. Flasks were individually sealed in an air-tight container along with an open vial containing 0.5 ml of 1 M NaOH. After 5 hr of incubation at 37 °C, the NaOH was neutralized with 0.5 ml of 1 M HCl and the amount of Na_2_^14^CO_3_ present was quantified via scintillation counting. Values were expressed as %CO_2_ release relative to the radioactive counts for the wild type.

### Membrane polarization, intracellular ATP determination and oxygen consumption rate

Membrane potential, ATP intracellular levels and oxygen consumption rate (OCR) were determined using the same culture settings. Mtb strains were cultured in modified Sauton’s medium until the mid-exponential phase (OD_580nm_ of 0.5-1). Bacteria were then centrifuged (4500 rpm; 10 min) and resuspended in fresh medium at a final volume 10 ml and a final OD_580nm_ of 0.5. Vehicle and OA were then added to the cultures at final concentration 250*μ*M and 500*μ*M. Samples were harvested after 24 hours treatment. For membrane potential, bacteria were resuspended in fresh medium with the same treatment conditions to a final OD_580nm_ of 1 and treated with 15 µM DiOC2 for 30 min at room temperature. Protonophore carbonyl-cyanide 3-chlorophenylhydrazone (CCCP) was added at a final concentration of 50 µM to provide a depolarized membrane control. Samples were then washed in fresh media with the same treatment conditions, and 200 µL in triplicate was transferred to a black clear bottom 96-well plates (Costar). Fluorescence measurements were performed in a SpectraMax M5 spectrofluorimeter (Molecular Devices): green fluorescence (488 nm/530 nm) and red fluorescence (488 nm/610 nm). Membrane potential was estimated as a ratio of red fluorescence over green fluorescence. ATP intracellular levels were determined using the commercial kit BacTiter-Glo (Promega) following the manufacturer’s instructions. To measure OCR, we used a Clark-type electrode system (Oxytherm+, Hansatech) with temperature set to 37 °C. Instrument calibration was performed following the manufacturer’s instructions. For OCR measurements, we added 1ml of culture and followed dissolved oxygen concentration over time the software Oxytrace+. OCR was calculated in the same software.

### Mouse infection

Mouse experiments were performed in accordance with the Guide for the Care and Use of Laboratory Animals of the National Institutes of Health, with approval from the Institutional Animal Care and Use Committee of Weill Cornell Medicine. Forty-eight female, eight-week-old Mus musculus C57BL/6 (Jackson Labs) were infected with ∼100–200 CFU/mouse through aerosolization using an Inhalation Exposure System (Glas-Col). Strains were grown to mid-exponential phase and single-cell suspensions were prepared in PBS with 0.05% Tween 80, and then resuspended in PBS to remove reagents. Lungs and spleen were homogenized in PBS + fatty acid free BSA (Roche) 0.05% to avoid LCFA growth inhibition in plate outgrowth. Homogenates were spread on solid modified Sauton’s medium to enumerate CFU (bacterial load). At day 70 post infection the small lung lobe was fixed with PFA 4% and stained (hematoxylin and eosin). Morphometric analysis of the lung pathology was performed as described elsewhere^61,62^ (2,3), with the software Interactive Learning and Segmentation Toolkit (Ilastik version 1.3.3). The probability maps of the whole lung and lesions were analyzed in CellProfiler Analyst (version 3.1.5). Lesion percentage was defined by the area occupied by lesion in each lung.

### Caseum surrogate

The surrogate caseum matrix was generated as previously described^9,63^, with minor modifications. Briefly, THP-1 monocytes were cultured in RPMI 1640 supplemented with GlutaMAX™, 10% fetal bovine serum (FBS) and 0.01 M Hepes. Cells were seeded at a density of 1 × 106/mL, and differentiated into macrophages with 100 nM phorbol 12-myristate 13-acetate (PMA), for 24 h. To induce intracellular lipid accumulation, the differentiated macrophages were exposed to 100 µM stearic acid for an additional 24 hours. Foamy macrophages were then detached using 5 mM EDTA and gentle scraping, washed three times with PBS, and pelleted by centrifugation at 300 × g for 5 minutes. Cell pellets were subjected to repeated freeze-thaw cycles to ensure cell lysis, followed by a 30-min incubation at 75 °C, resulting in the caseum surrogate. To assess bacterial survival, caseum surrogate pellets were dissolved in water and homogenized with 1.0 mm glass beads. Bacteria were grown to mid-exponential phase OD_580nm_ was adjusted to 0.6 before being added to the surrogate caseum at a final ratio of 2:1 (bacterial suspension volume:caseum weight). Bacterial viability was quantified through CFU enumeration.

### Statistical analysis

Generation of graphics and data analyses were performed in Prism version 9.0 software (GraphPad).

## DATA AVAILABILITY

TnSeq and whole genome sequencing raw data were deposited in NCBI’s Sequence Read Archive (SRA) under BioProjects PRJNA1247388 and PRJNA1248960.

## ACKNOWLEDGEMENTS

We thank Luming Chan, Frances Marks and Carolina Trujillo for support in cloning, sample processing and mouse aerosol infections. We thank Jansy Sarathy for technical advice in the implementation of the caseum surrogate model. This work was supported by the NIH grants R21 AI168506-01A1 and R01AI150916. The project leading to these results has received funding from “la Caixa” Foundation under the project code LCF/PR/HR24/52440021. Tiago Beites was supported by Fundação para a Ciência e Tecnologia with an individual grant (2022.00436.CEECIND). Marco Silva was supported by a Fundação para a Ciência e Tecnologia individual PhD fellowship (2023.00360.BD).

## REFERENCES

1 Global Tuberculosis Report 2024. (World Health Organization, 2024).

2 Peyron, P. et al. Foamy macrophages from tuberculous patients’ granulomas constitute a nutrient-rich reservoir for M. tuberculosis persistence. PLoS Pathog 4, e1000204 (2008). 10.1371/journal.ppat.1000204

3 Kim, M. J. et al. Caseation of human tuberculosis granulomas correlates with elevated host lipid metabolism. EMBO Mol Med 2, 258–274 (2010). 10.1002/emmm.201000079

4 Jaisinghani, N. et al. Necrosis Driven Triglyceride Synthesis Primes Macrophages for Inflammation During Mycobacterium tuberculosis Infection. Front Immunol 9, 1490 (2018). 10.3389/fimmu.2018.01490

5 Bedard, M. et al. A terpene nucleoside from M. tuberculosis induces lysosomal lipid storage in foamy macrophages. J Clin Invest 133 (2023). 10.1172/jci161944

6 Scriba, T. J., Maseeme, M., Young, C., Taylor, L. & Leslie, A. J. Immunopathology in human tuberculosis. Sci Immunol 9, eado5951 (2024). 10.1126/sciimmunol.ado5951

7 Sarathy, J. P. & Dartois, V. Caseum: a Niche for Mycobacterium tuberculosis Drug-Tolerant Persisters. Clin Microbiol Rev 33 (2020). 10.1128/cmr.00159-19

8 Sarathy, J. P. et al. Extreme Drug Tolerance of Mycobacterium tuberculosis in Caseum. Antimicrob Agents Chemother 62 (2018). 10.1128/aac.02266-17

9 Sarathy, J. P. et al. A Novel Tool to Identify Bactericidal Compounds against Vulnerable Targets in Drug-Tolerant M. tuberculosis found in Caseum. mBio 14, e0059823 (2023). 10.1128/mbio.00598-23

10 Bloch, H. & Segal, W. Biochemical differentiation of Mycobacterium tuberculosis grown in vivo and in vitro. J Bacteriol 72, 132–141 (1956). 10.1128/jb.72.2.132-141.1956

11 Daniel, J., Maamar, H., Deb, C., Sirakova, T. D. & Kolattukudy, P. E. Mycobacterium tuberculosis uses host triacylglycerol to accumulate lipid droplets and acquires a dormancy-like phenotype in lipid-loaded macrophages. PLoS Pathog 7, e1002093 (2011). 10.1371/journal.ppat.1002093

12 Lee, W., VanderVen, B. C., Fahey, R. J. & Russell, D. G. Intracellular Mycobacterium tuberculosis exploits host-derived fatty acids to limit metabolic stress. J Biol Chem 288, 6788–6800 (2013). 10.1074/jbc.M112.445056

13 Laval, T. et al. De novo synthesized polyunsaturated fatty acids operate as both host immunomodulators and nutrients for Mycobacterium tuberculosis. Elife 10 (2021). 10.7554/eLife.71946

14 Pandey, A. K. & Sassetti, C. M. Mycobacterial persistence requires the utilization of host cholesterol. Proc Natl Acad Sci U S A 105, 4376–4380 (2008). 10.1073/pnas.0711159105

15 Chang, J. C., Harik, N. S., Liao, R. P. & Sherman, D. R. Identification of Mycobacterial genes that alter growth and pathology in macrophages and in mice. J Infect Dis 196, 788–795 (2007). 10.1086/520089

16 Van der Geize, R. et al. A gene cluster encoding cholesterol catabolism in a soil actinomycete provides insight into Mycobacterium tuberculosis survival in macrophages. Proc Natl Acad Sci U S A 104, 1947–1952 (2007). 10.1073/pnas.0605728104

17 Nazarova, E. V. et al. Rv3723/LucA coordinates fatty acid and cholesterol uptake in Mycobacterium tuberculosis. Elife 6 (2017). 10.7554/eLife.26969

18 Arellano, H., Nardello-Rataj, V., Szunerits, S., Boukherroub, R. & Fameau, A. L. Saturated long chain fatty acids as possible natural alternative antibacterial agents: Opportunities and challenges. Adv Colloid Interface Sci 318, 102952 (2023). 10.1016/j.cis.2023.102952

19 Chamberlain, N. R. et al. Correlation of carotenoid production, decreased membrane fluidity, and resistance to oleic acid killing in Staphylococcus aureus 18Z. Infect Immun 59, 4332–4337 (1991). 10.1128/iai.59.12.4332-4337.1991

20 Greenway, D. L. & Dyke, K. G. Mechanism of the inhibitory action of linoleic acid on the growth of Staphylococcus aureus. J Gen Microbiol 115, 233–245 (1979). 10.1099/00221287-115-1-233

21 Won, S. R. et al. Oleic acid: an efficient inhibitor of glucosyltransferase. FEBS Lett 581, 4999–5002 (2007). 10.1016/j.febslet.2007.09.045

22 Kondo, E. & Kanai, K. The lethal effect of long-chain fatty acids on mycobacteria. Jpn J Med Sci Biol 25, 1–13 (1972). 10.7883/yoken1952.25.1

23 Beites, T. et al. Plasticity of the Mycobacterium tuberculosis respiratory chain and its impact on tuberculosis drug development. Nat Commun 10, 4970 (2019). 10.1038/s41467-019-12956-2

24 Beites, T. et al. Multiple acyl-CoA dehydrogenase deficiency kills Mycobacterium tuberculosis in vitro and during infection. Nat Commun 12, 6593 (2021). 10.1038/s41467-021-26941-1

25 Wong, A. I. et al. Cyclic AMP is a critical mediator of intrinsic drug resistance and fatty acid metabolism in M. tuberculosis. Elife 12 (2023). 10.7554/eLife.81177

26 Mortensen, J. E., Shryock, T. R. & Kapral, F. A. Modification of bactericidal fatty acids by an enzyme of Staphylococcus aureus. J Med Microbiol 36, 293–298 (1992). 10.1099/00222615-36-4-293

27 Kengmo Tchoupa, A. et al. Lipase-mediated detoxification of host-derived antimicrobial fatty acids by Staphylococcus aureus. Commun Biol 7, 572 (2024). 10.1038/s42003-024-06278-3

28 Banerjee, A. et al. A universal stress protein (USP) in mycobacteria binds cAMP. J Biol Chem 290, 12731–12743 (2015). 10.1074/jbc.M115.644856

29 Banerjee, A. et al. Cyclic AMP binding to a universal stress protein in Mycobacterium tuberculosis is essential for viability. J Biol Chem 300, 107287 (2024). 10.1016/j.jbc.2024.107287

30 DeJesus, M. A., Ambadipudi, C., Baker, R., Sassetti, C. & Ioerger, T. R. TRANSIT--A Software Tool for Himar1 TnSeq Analysis. PLoS Comput Biol 11, e1004401 (2015). 10.1371/journal.pcbi.1004401

31 DeJesus, M. A. et al. Comprehensive Essentiality Analysis of the Mycobacterium tuberculosis Genome via Saturating Transposon Mutagenesis. mBio 8 (2017). 10.1128/mBio.02133-16

32 Bosch, B. et al. Genome-wide gene expression tuning reveals diverse vulnerabilities of M. tuberculosis. Cell 184, 4579-4592.e4524 (2021). 10.1016/j.cell.2021.06.033

33 Lougheed, K. E., Thomson, M., Koziej, L. S., Larrouy-Maumus, G. & Williams, H. D. A universal stress protein acts as a metabolic rheostat controlling carbon flux in mycobacteria. bioRxiv, 2022.2006.2010.495724 (2022). 10.1101/2022.06.10.495724

34 Beg, M. A. et al. Mechanistic Insight into the Enzymatic Inhibition of β-Amyrin against Mycobacterial Rv1636: In Silico and In Vitro Approaches. Biology (Basel) 11 (2022). 10.3390/biology11081214

35 Beg, M. A. et al. Potential Efficacy of β-Amyrin Targeting Mycobacterial Universal Stress Protein by In Vitro and In Silico Approach. Molecules 27 (2022). 10.3390/molecules27144581

36 Chakraborti, S., Chakraborty, M., Bose, A., Srinivasan, N. & Visweswariah, S. S. Identification of Potential Binders of Mtb Universal Stress Protein (Rv1636) Through an in silico Approach and Insights Into Compound Selection for Experimental Validation. Front Mol Biosci 8, 599221 (2021). 10.3389/fmolb.2021.599221

37 Gould, T. A., van de Langemheen, H., Muñoz-Elías, E. J., McKinney, J. D. & Sacchettini, J. C. Dual role of isocitrate lyase 1 in the glyoxylate and methylcitrate cycles in Mycobacterium tuberculosis. Mol Microbiol 61, 940–947 (2006). 10.1111/j.1365-2958.2006.05297.x

38 Upton, A. M. & McKinney, J. D. Role of the methylcitrate cycle in propionate metabolism and detoxification in Mycobacterium smegmatis. Microbiology (Reading) 153, 3973–3982 (2007). 10.1099/mic.0.2007/011726-0

39 Eoh, H. & Rhee, K. Y. Methylcitrate cycle defines the bactericidal essentiality of isocitrate lyase for survival of Mycobacterium tuberculosis on fatty acids. Proc Natl Acad Sci U S A 111, 4976–4981 (2014). 10.1073/pnas.1400390111

40 Sweeney, M. I., Carranza, C. E. & Tobin, D. M. Understanding Mycobacterium tuberculosis through its genomic diversity and evolution. PLoS Pathog 21, e1012956 (2025). 10.1371/journal.ppat.1012956

41 Ehrt, S., Schnappinger, D. & Rhee, K. Y. Metabolic principles of persistence and pathogenicity in Mycobacterium tuberculosis. Nat Rev Microbiol 16, 496–507 (2018). 10.1038/s41579-018-0013-4

42 Sassetti, C. M., Boyd, D. H. & Rubin, E. J. Genes required for mycobacterial growth defined by high density mutagenesis. Mol Microbiol 48, 77–84 (2003). 10.1046/j.1365-2958.2003.03425.x

43 Griffin, J. E. et al. Cholesterol catabolism by Mycobacterium tuberculosis requires transcriptional and metabolic adaptations. Chem Biol 19, 218–227 (2012). 10.1016/j.chembiol.2011.12.016

44 Minato, Y. et al. Genomewide Assessment of Mycobacterium tuberculosis Conditionally Essential Metabolic Pathways. mSystems 4 (2019). 10.1128/mSystems.00070-19

45 Chugh, S. et al. Polyphosphate kinase-1 regulates bacterial and host metabolic pathways involved in pathogenesis of Mycobacterium tuberculosis. Proc Natl Acad Sci U S A 121, e2309664121 (2024). 10.1073/pnas.2309664121

46 Cole, S. T. et al. Deciphering the biology of Mycobacterium tuberculosis from the complete genome sequence. Nature 393, 537–544 (1998). 10.1038/31159

47 Drumm, J. E. et al. Mycobacterium tuberculosis universal stress protein Rv2623 regulates bacillary growth by ATP-Binding: requirement for establishing chronic persistent infection. PLoS Pathog 5, e1000460 (2009). 10.1371/journal.ppat.1000460

48 Glass, L. N. et al. Mycobacterium tuberculosis universal stress protein Rv2623 interacts with the putative ATP binding cassette (ABC) transporter Rv1747 to regulate mycobacterial growth. PLoS Pathog 13, e1006515 (2017). 10.1371/journal.ppat.1006515

49 Farewell, A., Diez, A. A., DiRusso, C. C. & Nyström, T. Role of the Escherichia coli FadR regulator in stasis survival and growth phase-dependent expression of the uspA, fad, and fab genes. J Bacteriol 178, 6443–6450 (1996). 10.1128/jb.178.22.6443-6450.1996

50 Yousuf, S., Angara, R., Vindal, V. & Ranjan, A. Rv0494 is a starvation-inducible, auto-regulatory FadR-like regulator from Mycobacterium tuberculosis. Microbiology (Reading) 161, 463–476 (2015). 10.1099/mic.0.000017

51 Ji, L. et al. Mycobacterium tuberculosis Rv0494 Protein Contributes to Mycobacterial Persistence. Infect Drug Resist 16, 4755–4762 (2023). 10.2147/idr.S419914

52 Aguilar-Ayala, D. A. et al. The transcriptome of Mycobacterium tuberculosis in a lipid-rich dormancy model through RNAseq analysis. Sci Rep 7, 17665 (2017). 10.1038/s41598-017-17751-x

53 Sheu, C. W. & Freese, E. Effects of fatty acids on growth and envelope proteins of Bacillus subtilis. J Bacteriol 111, 516–524 (1972). 10.1128/jb.111.2.516-524.1972

54 Parsons, J. B., Yao, J., Frank, M. W., Jackson, P. & Rock, C. O. Membrane disruption by antimicrobial fatty acids releases low-molecular-weight proteins from Staphylococcus aureus. J Bacteriol 194, 5294–5304 (2012). 10.1128/jb.00743-12

55 Kvint, K., Nachin, L., Diez, A. & Nyström, T. The bacterial universal stress protein: function and regulation. Curr Opin Microbiol 6, 140–145 (2003). 10.1016/s1369-5274(03)00025-0

56 Long, J. E. et al. Identifying essential genes in Mycobacterium tuberculosis by global phenotypic profiling. Methods Mol Biol 1279, 79–95 (2015). 10.1007/978-1-4939-2398-4_6

57 Andrews, S. FastQC: a quality control tool for high throughput sequence data., <http://www.bioinformatics.babraham.ac.uk/projects/fastqc> (2010).

58 Li, H. & Durbin, R. Fast and accurate long-read alignment with Burrows-Wheeler transform. Bioinformatics 26, 589–595 (2010). 10.1093/bioinformatics/btp698

59 Li, H. et al. The Sequence Alignment/Map format and SAMtools. Bioinformatics 25, 2078–2079 (2009). 10.1093/bioinformatics/btp352

60 DePristo, M. A. et al. A framework for variation discovery and genotyping using next-generation DNA sequencing data. Nat Genet 43, 491–498 (2011). 10.1038/ng.806

61 Fonseca, K. L. et al. Deficiency in the glycosyltransferase Gcnt1 increases susceptibility to tuberculosis through a mechanism involving neutrophils. Mucosal Immunol 13, 836–848 (2020). 10.1038/s41385-020-0277-7

62 Maceiras, A. R., Silvério, D., Gonçalves, R., Cardoso, M. S. & Saraiva, M. Infection with hypervirulent Mycobacterium tuberculosis triggers emergency myelopoiesis but not trained immunity. Front Immunol 14, 1211404 (2023). 10.3389/fimmu.2023.1211404

63 Xie, M., Osiecki, P., Rodriguez, S., Dartois, V. & Sarathy, J. A Physiologically Relevant In Vitro Model of Nonreplicating Persistent Mycobacterium tuberculosis in Caseum. Curr Protoc 5, e70118 (2025). 10.1002/cpz1.70118

