## Supplementary Information, Supplementary data 1 for "The *Mycobacterium tuberculosis* long-chain fatty acid resistome reveals the universal stress protein TB15.3 as essential for infection": Pinto and Silva 2025 Supplementary Information.pdf

**Supplementary figures 1-3**

**Supplementary Tables 1 and 2.**

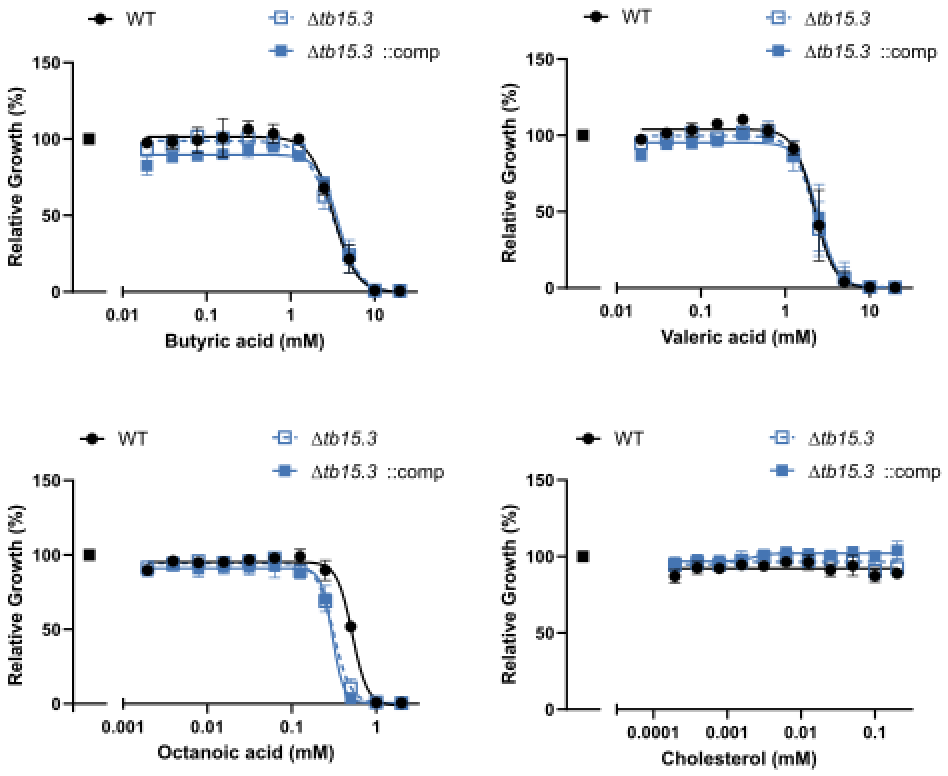


**Supplementary Fig. 1: Lipid susceptibility profiles.** Minimal inhibitory concentrations of fatty acids and cholesterol against Mtb wild type (WT), *tb15.3* knockout (Mtb Δ*tb15.3*) and complemented Mtb Δ*tb15.3* (Δ*tb15.3*::comp). Relative growth was calculated as $OD_{580nm}$ values at day 14 of lipid over vehicle control. Data are averages of three replicates and are representative of three independent experiments. Error bars correspond to standard deviation.

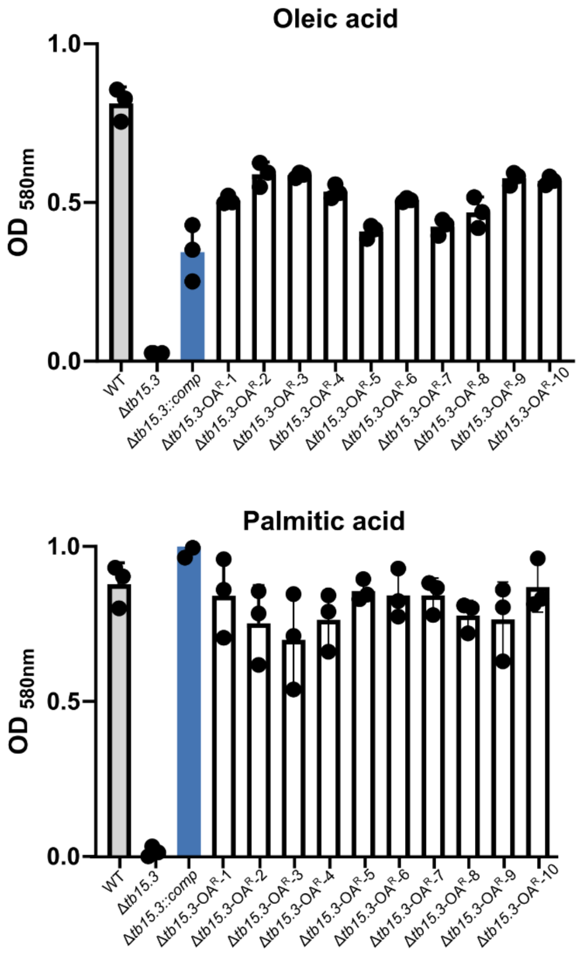


**Supplementary Fig. 2: OA resistance phenotype confirmation.** Mtb wild type (WT), *tb15.3* knockout (Mtb Δ*tb15.3*), complemented Mtb Δ*tb15.3* (Δ*tb15.3*::comp) and oleic acid resistant (OA$^{R}$) isolates were grown in modified Sauton's supplemented with oleic acid 500 µM or palmitic acid 250 µM. OD$_{580nm}$ was recorded after 21 days of culture. Data are averages of three replicates and are representative of two independent experiments. Error bars correspond to standard deviation. Data are averages of three replicates and are representative of three independent experiments. Error bars correspond to standard deviation.

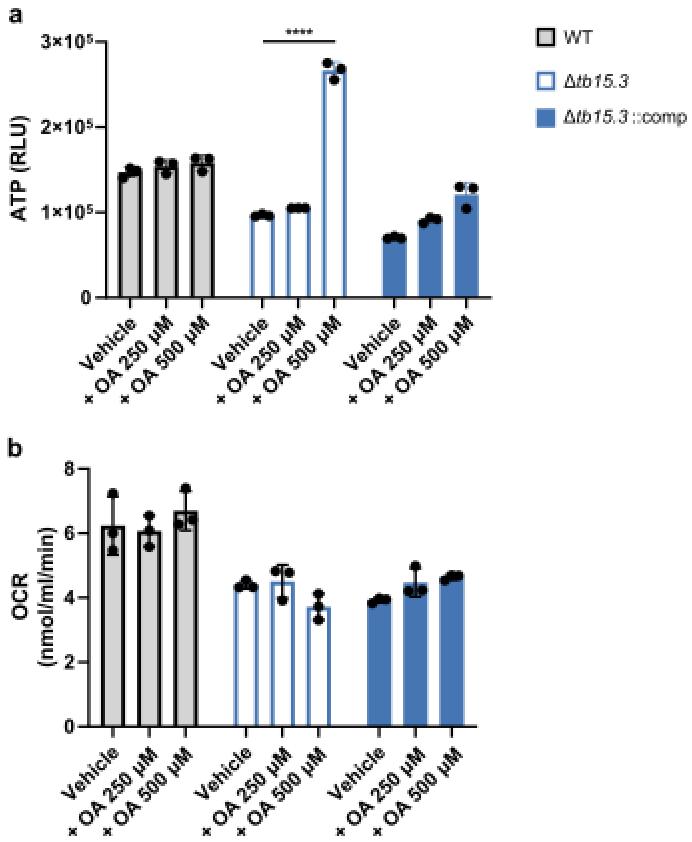


**Supplementary Fig. 3: Intracellular ATP levels (a) and oxygen consumption rate (OCR) (b) in response to oleic acid (OA).** Mtb wild type (WT), *tb15.3* knockout (Mtb Δ*tb15.3*) and complemented Mtb Δ*tb15.3* (Δ*tb15.3*::comp) were cultured in modified Sauton's and treated with OA 250 μM and 500 μM or vehicle control for 24 hours. Data are averages of three replicates and are representative of three independent experiments. Error bars correspond to standard deviation. Statistical significance was assessed by one-way ANOVA followed by post hoc test (Tukey test; GraphPad Prism). ****$P < 0.0001$

**Supplementary Table 1.** Additional polymorphisms identified in Δ*tb15.3* in comparison with parental strain.

| Gene | Position | Ref | Alt | #Ref reads | #Alt reads | AA change | Mutation |
|---|---|---|---|---|---|---|---|
| *PE_PGRS1* | 132415 | C | G | 0 | 121 | R346G | Missense |
| *lprO* | 210094 | A | G | 0 | 143 | NA | Silent |
| *PE_PGRS9* | 837036 | A | G | 0 | 81 | T445A | Missense |
| *Rv0887c;Rv 0888* | 986207 | C | A | 0 | 145 | (intergenic) | NA |
| *pknD* | 1039376 | C | G | 0 | 127 | M181I | Missense |
| *snoP* | 2933205 | G | T | 0 | 99 | D151E | Missense |
| *Rv2813; Rv2814c* | 3121725 | GGGTTTCCGT CCCCTCTCGG GGTTTTGGGT CTGACGACAT GCTGAGCTGA GGCGCCGGAT GATGGTGGTG CTGAA | G | 0 | 127 | (intergenic) | NA |
| *PPE46* | 3381807 | G | GC | 0 | 12 | NA | Frameshift Insert |
| *Rv2553c* | 2874461 | G | GCCA | 1 | 81 | NA | In-frame Insert |

Ref – reference sequence
Alt – aleternate sequence

**Supplementary Table 2.** Drug susceptibility profiles from 3 independent replicates.

| Compound | Target process | MIC (μg/ml) | | | | | |
|---|---|---|---|---|---|---|---|
| | | WT FA Free | WT + OA 100 μM | Δ*tb15.3* FA Free | Δ*tb15.3* + OA 100 μM | Δ*tb15.3*::comp FA Free | Δ*tb15.3*::comp + OA 100 μM |
| DDD00853663 | Oxidative phosphorylation | >10 | 0.59-1.21 | >10 | 0.09-0.13 | >10 | 0.28-0.54 |
| Clofazimine | | 0.91-2.46 | 0.8-2.29 | 0.88-1.10 | 0.28-0.57 | 0.82-1.38 | 1.10-1.27 |
| Q203 | | 0.01-0.03 | 0.01-0.03 | 0.01-0.03 | $2.45x10^{-4}$ - $3.85x10^{-8}$ | 0.01-0.02 | 0.01-0.02 |
| Bedaquiline | | 0.34-0.81 | 0.30-0.46 | 0.57-0.71 | 0.07-0.13 | 0.51-0.82 | 0.39-0.43 |
| Ethambutol | Arabinogalactan biosynthesis | 1.62-5.65 | 1.48-4.95 | 1.04-5.13 | 0.69-3.47 | 1.31-5.03 | 1.49-5.05 |
| Isoniazid | Mycolic acid biosynthesis | 0.01-0.02 | 0.02-0.04 | 0.01 | 0.01-0.02 | 0.01 | 0.02 |
| Rifampicin | Transcription | 0.63-1.6 | 0.44-1.48 | 0.57-2.15 | 0.11-0.64 | 0.78-2.29 | 0.63-1.75 |
| Amikacin | Translation | 0.35-0.58 | 0.47-0.69 | 0.36-0.52 | 0.18-0.37 | 0.33-0.49 | 0.61-1.14 |
| Linezolid | | 0.77-1.41 | 0.76-1.33 | 0.68-1.17 | 0.24-0.58 | 0.78-1.19 | 0.69-1.26 |
| Vancomycin | Peptidoglycan biosynthesis | 25.23-27.5 | 6.62-9.91 | 15.94-62.92 | 2.08-56.99 | 17.13-102.2 | 15.88-17.84 |
| Moxifloxacin | DNA replication | 0.1-0.16 | 0.08-0.15 | 0.08-0.22 | 0.04-0.14 | 0.09-0.16 | 0.13-0.20 |
